# The Ratchet Model of Synovial Flares in Palindromic Rheumatism

**DOI:** 10.1101/2024.12.06.626783

**Authors:** S. Chakraborty, S. Phatak, S. Rath, P. Goel

## Abstract

Synovial flares in palindromic rheumatism (PR) are aperiodic bursts of inflammation in the joints, which usually self-resolve in a timescale hours or days. PR patients are believed to transit to a chronic auto-immune disease called rheumatoid arthritis (RA) in most cases, however, many patients remain palindromic indefinitely. We utilize and adapt a minimal ODE model of rheumatoid arthritis (RA) developed by Baker et al. to study PR in greater detail. We address questions characterizing the incidence, decay and sustenance of synovial flares in palindromic patients. A key question is to describe the nature of the transition from palindromic to full RA. We show that PR flares ordinarily resolve spontaneously, however, there is a secondary equilibrium in the model into which the trajectory can sometimes get trapped. When this “meta-stable locking” occurs, it initiates an adaptation that helps rescue the flare. Furthermore, this adaptation in turn activates a secondary adaptation in response to fluctuations in the healthy steady state. Finally, we show that if metastable locking occurs frequently enough these adaptation sequences turn maladaptive and the system slowly progresses into fully developed RA.

## 1 Introduction

Palindromic rheumatism (PR) is a syndrome characterized by arthritic swelling, pain, and erythema in or around the synovium of the joints over brief periods. These inflammatory “flares” happen at varying intervals and typically resolve on their own. It is not clear whether they result in cellular changes in the synovial membrane which are temporary or persistent. The flares in PR occur on the peri-articular soft bone tissues but leave no radiographic lesions or residual damage, unlike in rheumatoid arthritis (RA) [1, 2]. A flare usually lasts for a few hours to days, while the syndrome itself can continue to recur for years. The lack of warning signs between inflammatory episodes complicates the prediction of subsequent bouts.

We aim to study a minimal mathematical model describing a synovial flare in PR in order to understand not only the self-resolving nature of the inflammation but also the relationship between one bout of inflammation and the next. Since synovial flares are thought to be triggered by external stimuli – which continue to advance and eventually recede spontaneously – the mathematical concept of excitable systems appears to be ideally suited to this aim. [3, 4, 5, 6] One such model is due to Baker et al. [7], in which the authors a develop a two-dimensional system of pro- and anti-inflammatory cytokines underlying rheumatoid arthritis. We note that in this study we are specifically interested in PR alone, rather than RA. Indeed, PR does finds a brief mention in the Baker model as a transitory intermediate between an age-related progression from a healthy to diseased state, as depicted in Fig. 1.

**Figure 1:**
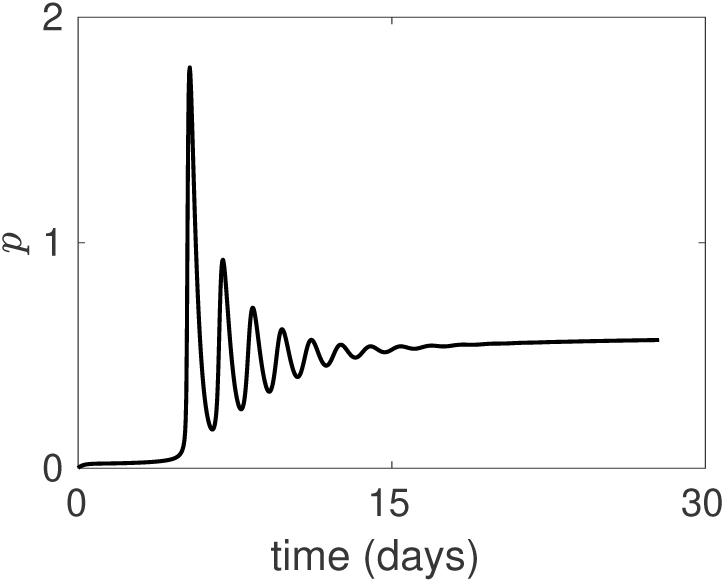
The palindromic state is characterized in Baker et al. [7] (see Fig. 12; reproduced here) as an intermittent stage that exists while a subject transitions from a healthy to a diseased stage. The oscillatory dynamics of the flares is actually damped, riding on top of a baseline inflammation that elevates steadily. Notice the first couple of flares are especially pronounced; this does not find evidence in the clinical literature. The entire transition to RA is seen take place within days; clinically, however, each individual flare can last several days. Parameters are as described in Section 2, together with 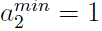, 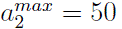, as in [7]; note, however, that we take 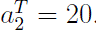.

The authors investigate several parameter regimes that lead to multistability and various bifurcations scenarios; in that language, PR occurs during a passage from one attractor state (healthy) to another (disease) under a relapsing-remitting framework. While this is an interesting explanation, we find that it does not explain a number of clinically observed features of PR flaring. In contrast to the Baker picture above, palindromic flares occur suddenly as unanticipated inflammatory episodes [8], independent of an agerelated decline [9]. In other words, synovial flares are expected to occur at irregular time intervals. Further, PR may not necessarily be a precursor to RA [10, 11, 12]. We enumerate a number of differences between the Baker view of PR flares and their clinical characteristics in Table 1.

**Table 1:** Differences of PR representation.

| Baker model | Clinical observations |
| --- | --- |
| PR is shown to be a periodic solution of $P$ and $A$ of constant frequency with short intra-width interval and same interval between flares. | PR is observed as an abrupt state of synovial flares, with varied inter-flare interval among patients. |
| PR is an intermediate stage to overt RA. | Clinically, two classes of phenotypes are observed: (I) Patient has small and infrequent number of flares, which consistently subside back to "normal" state i.e. patient remains palindromic. (II) Patient has entered chronic diseased state. |
| The timescale involved in the flares to reach the two steady states has been shown in days. | The transition from palindromic to RA typically takes years. The number can vary between years, even lasting a decade.[1, 11, 12] |

We aim to develop a description of PR using the Baker model as a template, however, we shift focus onto the clinical features of flares. In this paper *we emphasize that PR flares are better understood as excitable trajectories* instead. In fact, we retain much of the underlying dynamics and parameter regimes obtained in [7] but describe flares in terms of trajectories arising out of perturbations to the attractor states, which return to them. In this paper, we will show that this shift in perspective allows us to describe PR flares in line with clinical expectations.

The second question relates to the nature of life course changes that underlie progression of PR. On the one hand, some PR patients may never progress to persistent RA; in others, flares may serve as a prognostic warning of impending RA. In this paper we propose a **Ratchet model** of PR. Each flare episode can be thought of as leaving behind a subtle, largely unobserved residual defect in its wake. The key question is whether each PR flare resolves completely, or only apparently so; can the accumulation of a number of these result in progression of the disease? We aim to describe this picture of longitudinal changes within the framework developed here and hope to clarify related causal questions.

The paper is organized as follows. In Section 2, we describe the specifics of the Baker model. In Section 3, we describe excitability as a qualitative representation of a synovial flare and we show the region space in which it may exist. In Section 4, we further ask: what is the reason behind the longitudinal transition happening from a palindromic state to chronic disease? The “ratchet” model for disease progression is described in greater detail. We ask what can determine disease progression and its relation to the pathological changes as a result of a flare. We close with discussion in Section 5.

## 2 Description of the Model

The Baker model represents pro– and anti-inflammatory cytokines coupled in an activator-inhibitor motif, Figure 2. The model is composed of five fluxes driving the evolution of pro– and anti-inflammatory cytokines to healthy or disease states.

**Figure 2:**
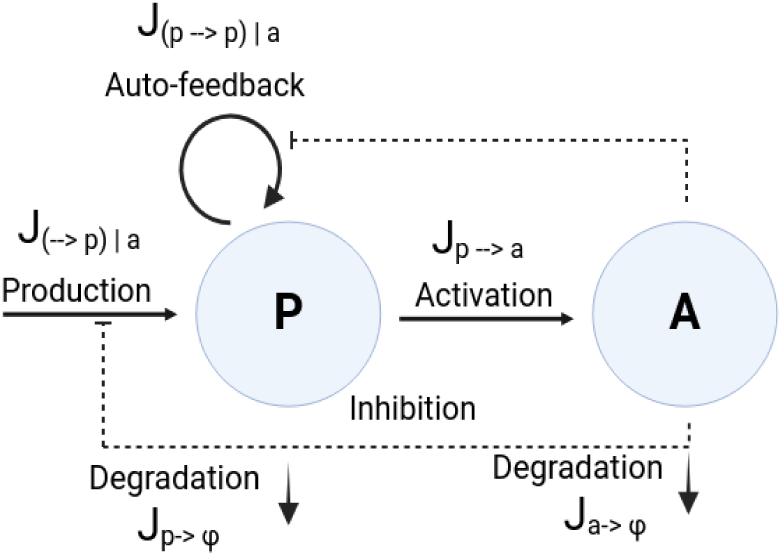
Pro-inflammatory cytokines *P* activate the production of anti-cytokines *A*, denoted by *J_p→a_*. *P* is produced by its constant background flux *J_→p|a_* along with its auto-positive feedback *J_p→p|a_*, both inhibited by *A*. The cytokines undergo linear degradation rates, with fluxes represented as *J_p→ϕ_* and *J_a→ϕ_*.

The equations of the model are:

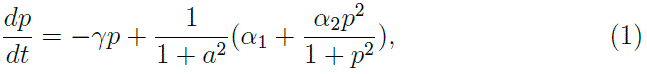

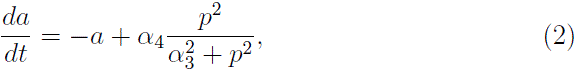

with typical parameters taken to be *α*_1_ = 0.025*, α*_2_ = 8*, α*_3_ = 0.5*, α*_4_ = 3.5*, γ* = 1.25. Here *α*_1_ represents the background production rate for proinflammatory cytokines, *α*_2_ represents the strength of pro-inflammatory auto feedback flux, *α*_3_ is the pro-inflammatory concentration at which anti-inflammatory concentration has reached half of it’s maximum value, *α*_4_ represents the strength of the production of anti-cytokines, and *γ* is the clearance ratio for the two classes of cytokines.

We note that *p* and *a* variables were obtained by nondimensionalising using arbitrary constants *c*_2_ and *c*_4_, while ‘t’ is made to be unitless by multiplying it with *d_a_* or degradation rate of anti-inflammatory cytokines; see [7] for more details.

## 3 Palindromic flares as excited trajectories

We view a PR synovial flare as an excitable event that starts and recedes by itself to the same *P* and *A* levels.

Here, we note, that the PR flare returns to the same equilibrium with no implication of systemic damage, as clinically observed [8, 13]. The model is capable of presenting transiently sensitized events, reflecting the influence of unknown triggers in a diseased system. The phase space shows two stable steady states, which suggests that the system rests in two different cytokine levels for prolonged periods of time. With certain perturbations, we show an observable amplification of the cytokines, and subsequent decay. However, niched perturbations transports the cytokine system to a “locked” state.

We use the two-parameter bifurcation structure of *α*_1_−*α*_2_ is to identify region spaces for bi- and monostability in the model. A cusp bifurcation emerges as the stable steady states annihilate with the saddle node to form a single dominating stable state. Our findings indicate that a small portion of bistable space exhibits excitable flare regimes, whereas the entire monostable area has the ability to produce flares. This suggests that the physiologal response in a palindromic patient is limited to an optimal range for the strength in autofeedback loop of the cytokines.

We describe the flare with its individual fluxes in Appendix 6.1. In Appendix 6.2 we use two parameter sets corresponding to 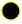 and 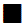 to compare and contrast flares in the bistable and monostable regimes. We demonstrate that although the flares appear similar in either situation, the phase planes corresponding to these regimes [7] are considerably different. In particular, we show that the inhibitory flux is weak in the bistable case, but strong in the monostable regime as detailed in Appendix 6.3.1.

## 4 The Ratchet model of PR progression

It is of considerable interest to determine whether PR flares represent an acute event or are indicative of a progression that ultimately results in RA. Several studies suggest there is no radiological damage that occurs to a PR patient following a flaring episode [1, 2]. It is often described an an “abortive” form of RA, which leaves no trace [14]. On the other hand, there is a substantial proportion of PR patients who do transition to RA over a period of time [1, 15].

There is, however, an underlying difficulty in characterising this progression. We need to address whether (i) each flare is somehow *responsible* for the degeneration, which in turn accumulates? Perhaps there is some shift in the cytokines’ state as a result of a flare. However, the return to equilibrium in the model suggests that this is not so. On the other hand, perhaps (ii) the flare is a *passive, symptomatic* event, and the underlying causal degenerative process is something else? In that case, what is its relationship to the flare, that is, how does it alter the flare?

We next describe a “ratchet model of PR”, which alludes to a progressively worsening disease state, that is, degenerative processes tend to accumulate with successive flares. Phenotypically, if we were to compare two flares, cytokine levels will be elevated following the latter. It can roughly become a marker, in a palindromic patient, progressing towards disease. However, there is a need to mechanistically pin down the reason for the progression.

We first note that a naive model is to simply assume there is a slow increase in the inflow of immune cells into the synovium; this can be expected to eventually leads to a rise in cytokine levels over a period of time. Figure 5 shows a simulation in which *α*_1_ is driven upwards, that is,

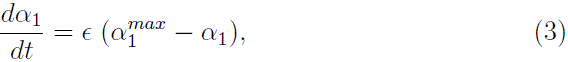

describes a straightforward way in which transition may occur. Intuitively, we might even obtain a similar outcome with a gradual increase feedback strength over time, that is, with *α*_2_. On the other hand, such an explanation does not provide us an insight about what might trigger the rise in *α*_1_ or *α*_2_.

There is, however, another viewpoint which is interesting to explore. Notice that due to the bistability it is possible that the trajectory can get locked in a metastable state as it attempts to return to its baseline origin (Fig. 3b). We argue, that a palindromic pathophysiology, which is primed to end up in chronic disease, will encounter more *frequent* locks throughout their life course. In addition to that, we understand that *A*’s production is solely driven by *P*, and thus results in low physiological levels of *A* as well at low values of *α*_1_. That would in turn, indicate a low coupling of the systemic cytokines to the synovium. Thus, we argue that, there must be a high inhibitory strength of anti-inflammatory cytokines coupled to control pro-cytokines to low background levels. It is expected to exist in the palindromic state, which prevents it from attaining the chronic state of the disease, during the initial phase of synovial attacks.

**Figure 3:**
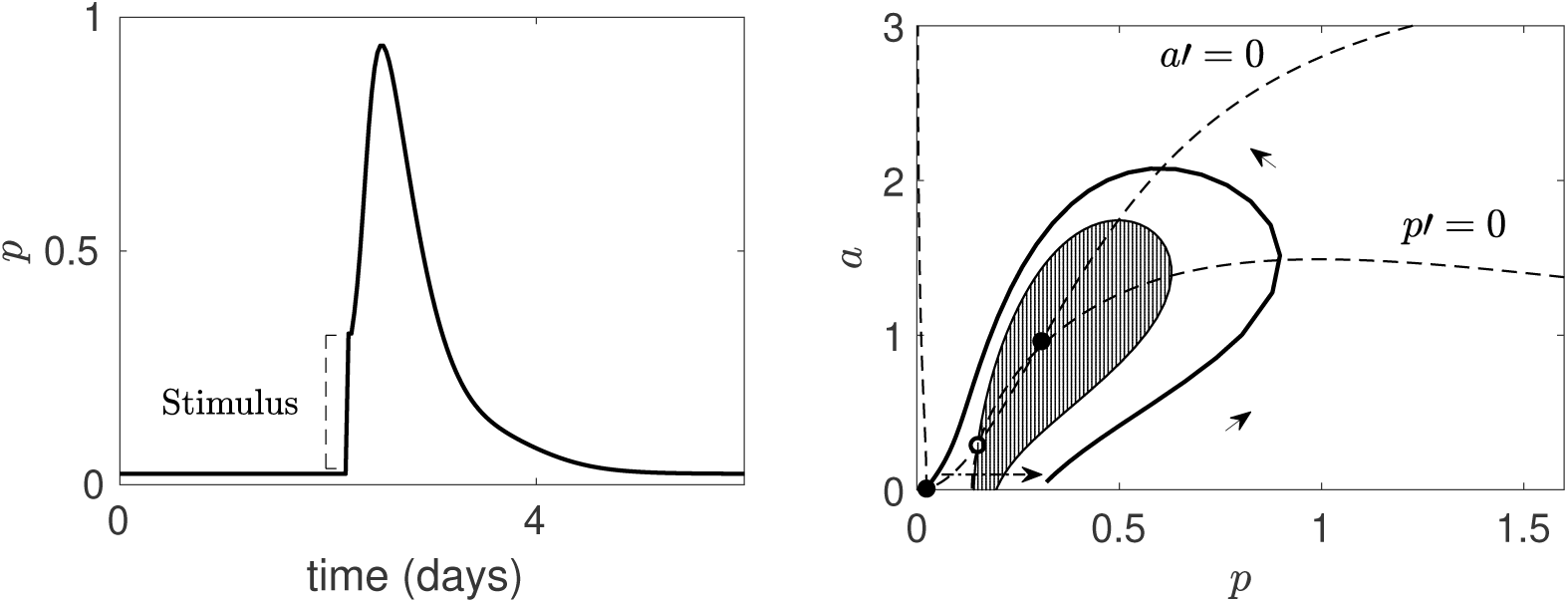
(A) A short stimulus is excitable and resembles a synovial flare of 1-1.5 days. (B) Phase-plane of *P*-*A*: Divides region flare (unshaded) and non-flare (shaded) regions. The system has two stable steady states (low, high values of cytokines) separated by an unstable state. Parameters are as described in Section 2.

**Figure 4:**
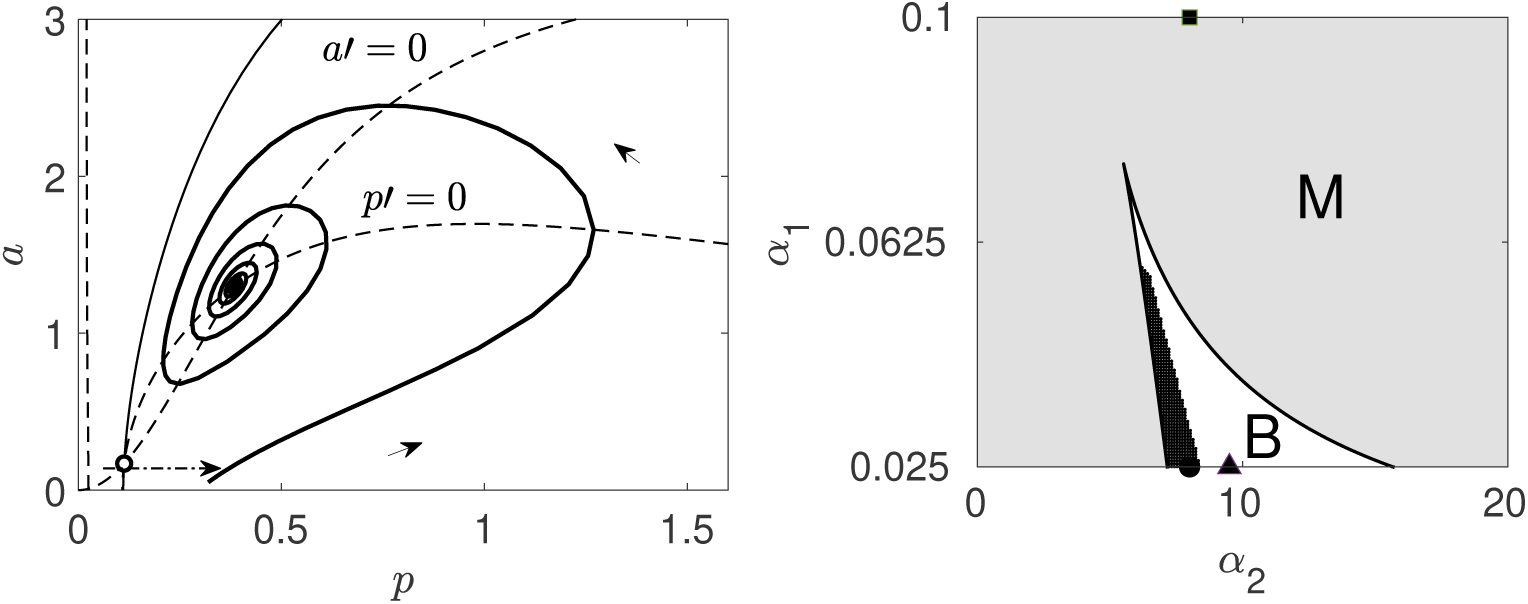
(A) Represents a phase-plane visualization of a flare that does not return to the same equilibrium. (B) Two-bifurcation diagram: Parameter combinations that *lead to a flare* are indicated by the shaded (dark) region inside “B” (bistable and the shaded (bright) region inside “M”/monostable. The triangle represents Fig.(4a), and the combinations of the white area in “B” (*α*_2_=9.5, parameters are described in Section [2]), the square represents a monostable flare in Fig.(6.2), and the black circle represents the flare in Fig.(3a).

**Figure 5:**
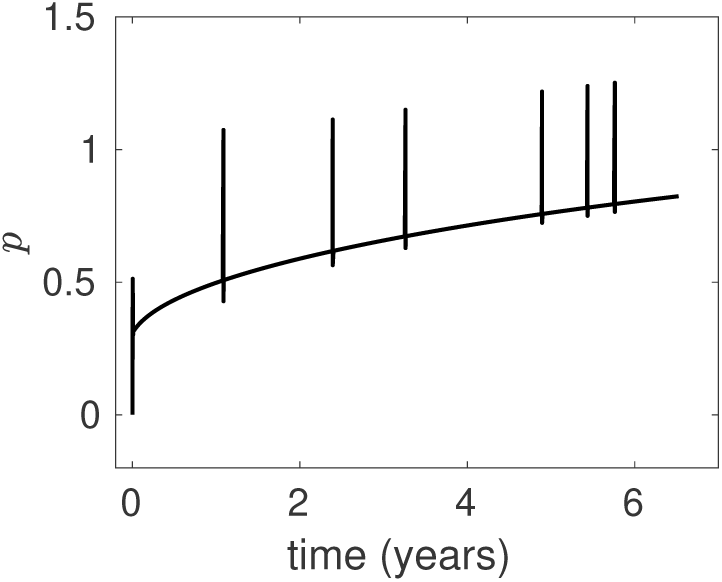
Each flare occurs on a fast timescale, while the influx of immune cells into the synovium (*α*_1_), which will eventually lead to an explosion of *p*, occurs on a very slow timescale. It represents a *ratcheting* form of progression, where the rise is occurring in a steady and irreversible manner. The flares usually occur aperiodically, depending on the incidence of triggers in the system.(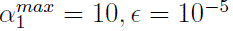, other parameters as described in Section 2)

**Figure 6:**
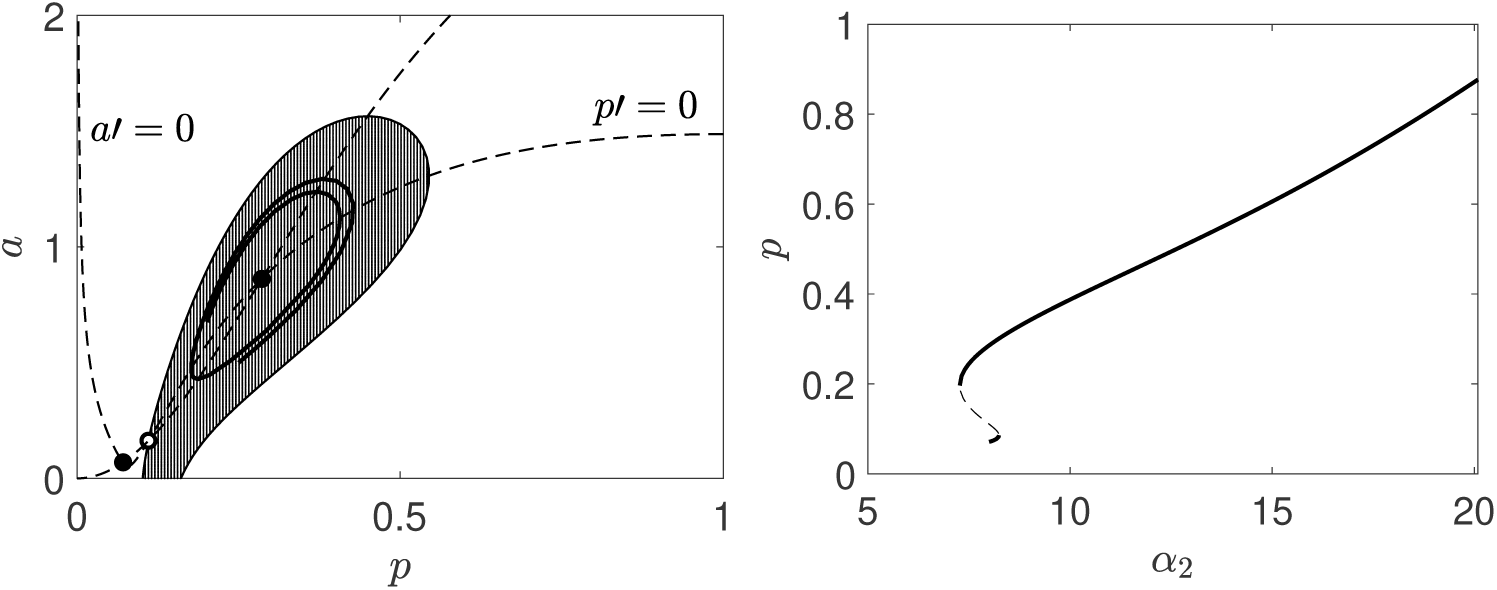
(left) The black trajectory is locked in the metastable region. We observe that the bistable structure is maintained, as from the original model. (*α*_1_=0.05, *f* =5; other parameters are s described in Section 2) The shaded region essentially covers a short space of *P*-*A*, so the locked stage might be rare. (right) The modified model shows a much lower range of *P* values, unto which the palindromic flare returns. Intuitively, it infers that the PR regime *can be possibly maintained* only due to a short range of strength of auto-feedback. A sudden sharp increase in *α*_2_ will drive the system to disease stage.

### 4.1 Strong inhibition of the production of ***P*** by ***A***

We show in Appendix 6.3.1 that the Baker model, as it stands, has weak inhibition on the production rate of *P* by *A*. This poor feedback results in a negligible influence of increasing *A* on (the production of) *P*. In other words, the feedback of *A* is limited to its modulation of the positive feedback auto-production of *P* but not its extrinsic production. Here we propose to modify this feature, that is, we argue that *A* is capable of exerting a strong influence on the rate at which *P* is produced to begin with. The dimensional version of the equations in the Baker model [7] are as follows:

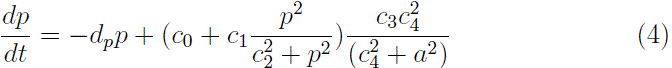

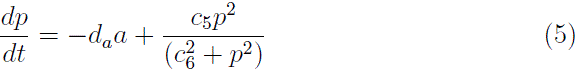

The Baker model [7] had used *c*_4_ to non-dimensionalise *A*, to reach the final set of equations. The physical interpretation of inhibition is accounted by the concentration of *A* in the system, and we have no way to account for the sensitivity of inhibition. In order to account for the *influence* of *A* on the constant flux, i.e. *α*_1_, we non-dimensionalise *A* by *c*_6_, to create an additional parameter *f*. Given that *α*_1_ bridges the connect from systemic levels of lymphocytes and inflammatory mediators to the synovium, we find it useful to use *f* to explain the effectiveness of inhibition by *A* on *P*. The flux analysis reveals that *f* has a bigger impact on the inhibitory flux rate, rather than *α*_4_ or *α*_3_.(6.4) Thus equation (2) is the same but equation (1) is modified to

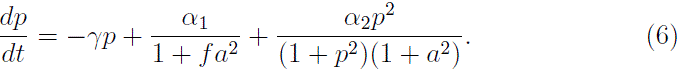

In the following section we study the behaviour of the stronger model. Our version of the model explains a tightly coupled inhibitory link from systemic inflammation to localized inflammation. The flaring of the synovium is a local event, and is mainly countered by the density of *A* produced and the respective decay rates. While the continuous influx of immune cells aggravates with continued insults and degrading pathophysiology [12, 16], the modulatory strength of inhibition would be physiologically relevant in explaining immune self-tolerance.

### 4.2 Adaptation of *α*_4_ releases metastable locking

We have observed potential triggers that led to a phenotypical flare, that can be observed and investigated clinically. However, we also found a metastable region in the palindromic phase-space, wherein certain perturbations would get locked at higher values of cytokines.

We now introduce an adaptation. Potentiating *α*_4_ is the key step required to release any metastable locking of the flare. Once *α*_4_ is high, it will lead to the production of more anti-cytokines in the system. This moves the system out of the lock, and once it does, *α*_4_ returns to its normal levels. Thus, the system’s adaptation to these bursts or spikes in *P* engages a protective response to resolve it.

We define the following equations:

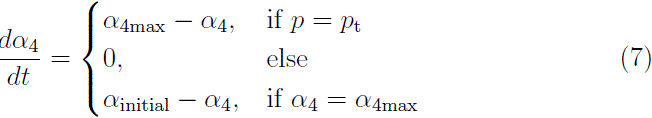

Here, *p*_t_ defines an arbitrary threshold for the metastable state. Once the pro-cytokines cross the threshold, the adaptation system is triggered.

Fig. 7a illustrates a locked state in *P*, which oscillates to the higher equilibrium, which get’s resolved as *α*_4_ (Fig. 7d) evolves. Thus, this system is capable of responding to sustained elevation of cytokine levels. Consequently, we show that the simulations that span several years in Fig.(7b, 7c, 7e, 7f) depict a realistic progression of palindromic physiology. Notably, the base equilibrium in between flares remains unchanged and is characteristic of the pathophysiology remaining palindromic. We present cases where locked stages of inflammation are interspersed with distinct synovial flares. The key insight is that palindromic systems are likely to undergo *more locked states*, which can predispose them to chronic disease. However, as we demonstrate in the next section, the restoration process can occasionally become harmful due to secondary responses within the adaptive system.

**Figure 7:**
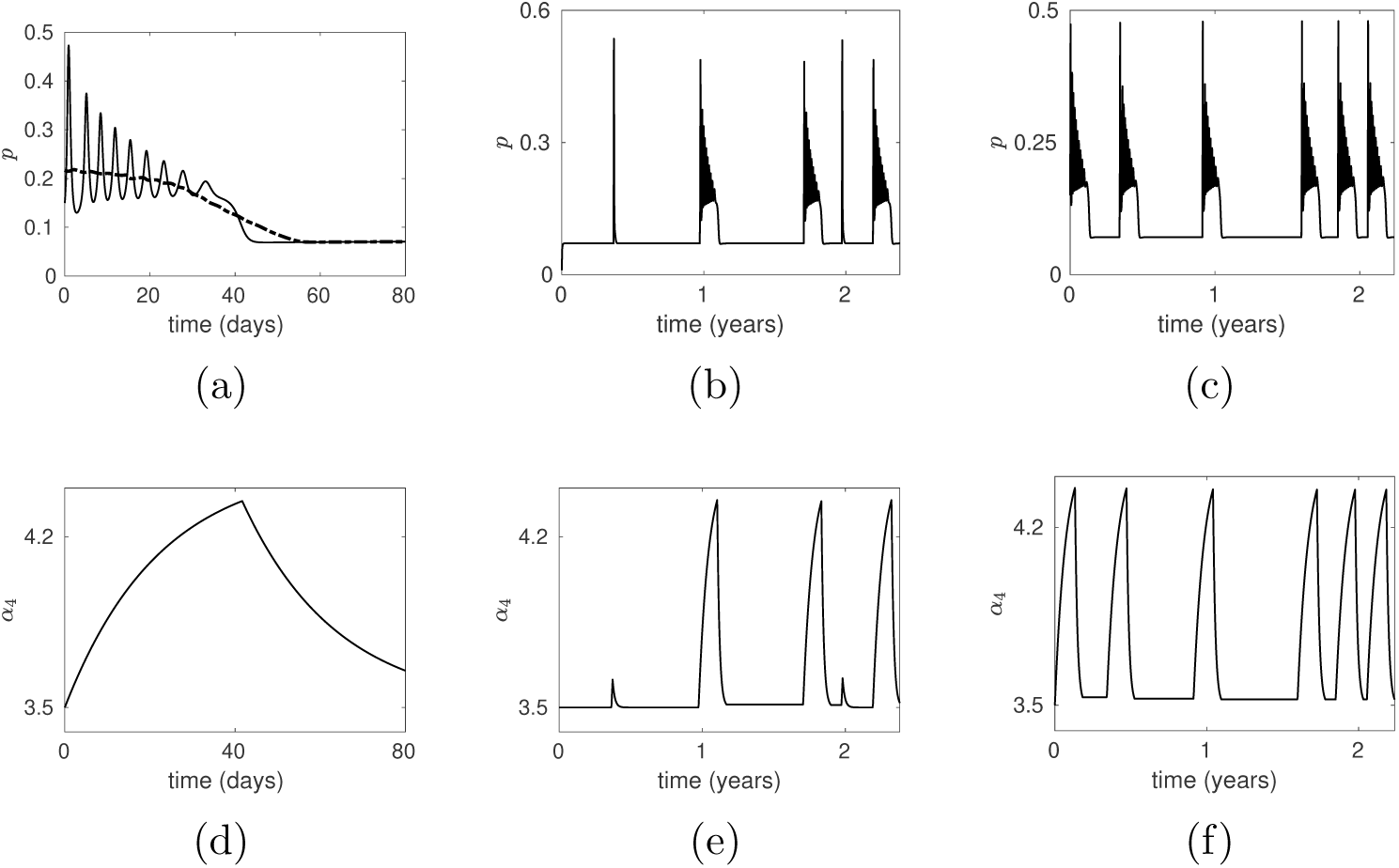
(a) *P* is oscillating steadily to the metastable state, but relaxes to the lower steady state after a while. (d) On sensing the lock of the system sets in, *α*_4_ is potentiated, and also falls subsequently. (b) and (e) show synovial flares, with intermittent insults, recovered by *α*_4_. (c) and (f) show multiple *aperiodic* insults; all of the above events are occurring on a timescale of years. (*α*_4_*_max_* = 4.5*, p_t_* = 0.1, other parameters as described in Section 2)

### 4.3 Multiple locked-state insults lead to a secondary response in ***f***

We note that changes in *α*_4_ also initiate changes in the *physiologically healthy levels of cytokines* in the system. It is plausible that *P* ought to be maintained homeostatically at low baseline levels in healthy individuals, for multiple processes like response to inflammogens, wound repair, cell differentiation, regulation of the innate and adaptive immune response and so on [17, 18]. A significant drop in physiological levels of *P* is likely to be detrimental to the system. Thus, a compensatory mechanism is initiated. We thus argue that changes in *P* which occur after each bout of locked flares, would initiate a secondary response.

Fig. 8a shows that the *a*-nullcline “wobbles” whenever a metastable lock and its subsequent *α*_4_ adaptation occur. This is turn causes the *p* corresponding to the healthy steady state to get slightly depressed, albeit temporarily. One way in which the system might react to this *p*-lowering is to try to increase the production flux of *p*. This can be achieved by coupling changes in *α*_4_ to the incoming flux, that is, by having *f* in Eq. 6 decrease whenever *α*_4_ changes from baseline (from below). In other words:

1. *α*_4_ rises in response to a metastable lock and releases it.
2. This, however, causes suppression of the *p*-cytokine in the healthy state. We note that this is a latent state in the system during this time; the trajectory itself is an elevated state while returning to the healthy fixed point. Nonetheless, we argue that the system senses this deviation^1^ and
3. simultaneously triggers a suppression in *f* into order to effectively increases the effective *α*_1_ flux (see Eq. 6) and drive *p* in.

**Figure 8:**
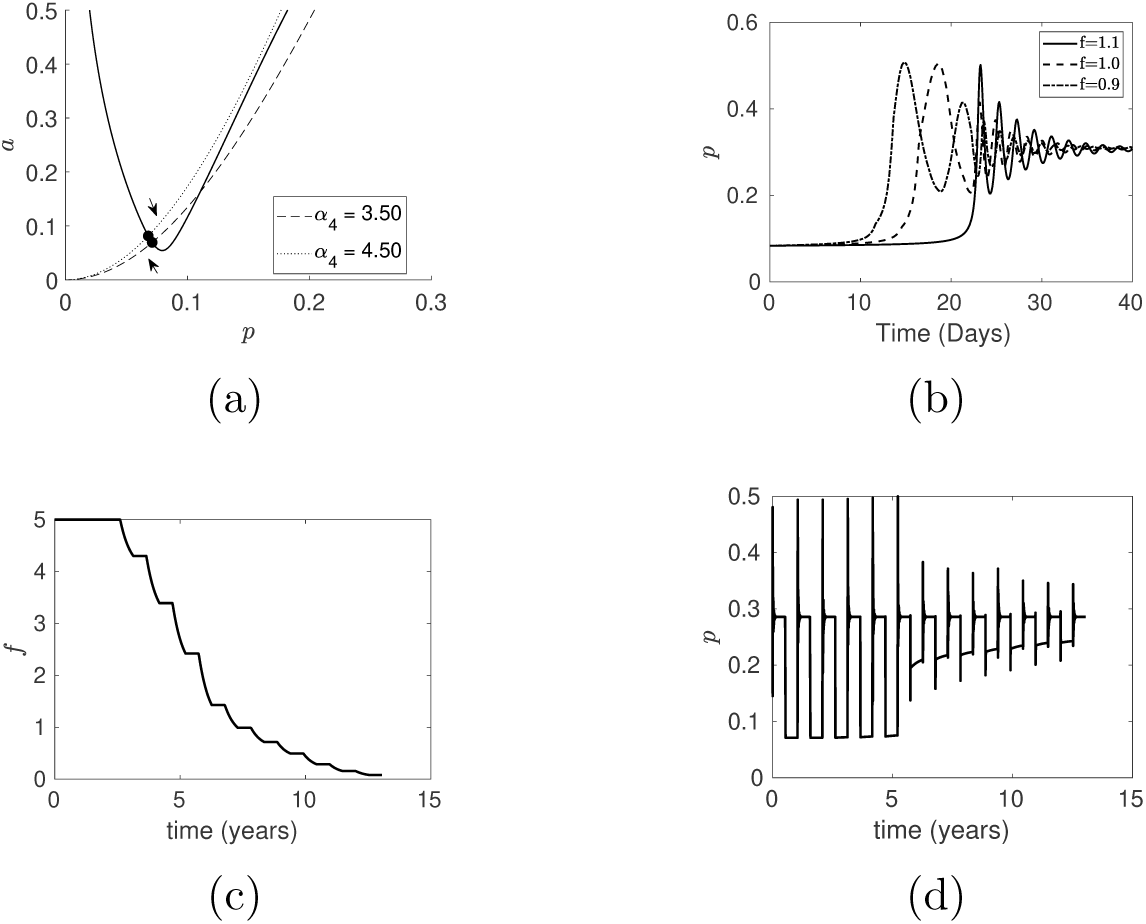
(a) Unsteady changes in the physiological levels of *P*. (b) Patients near the bifurcation point, take longer time to get locked. (c) We have represented a decrease in the inhibitory effectiveness as a stepwise manner, over the years (d) Consequent changes in *P* occurs, and shows a sharp transition over a critical *f* value. (*f_crit_* = 1.1, *ɛ* = 2.5 ∗ 10*^−^*^4^, other parameters as described in Fig. 7f and Section 2)

We use the following to model *f* in response to *α*_4_.

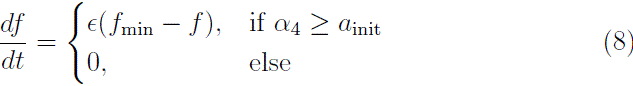

where, *ɛ* controls how fast *f* decays during the *α*_4_ bout.

Fig. 8a shows the “wobble” effect in cytokines, as the *α*_4_ dynamics unfold over multiple locking stages. We show subsequent decrease in the levels of *f* over time, in response to *α*_4_ bursts(Fig. 8c). The system is sensing fluctuations in the lower levels of *P*, which may interfere with critical physiological processes. At a particular *f_crit_* the system can take longer for the system to get locked in the metastable state, thus lead to longer time in transition over the years. It describes the situation where the physiological system is close to the “ghost” region of the saddle node bifurcation, where the vector fields are the slowest(Fig. 8b). It introduces variability among patients, with some remaining affected for significantly longer periods. It is important to note that the *qualitative* nature of the transition does not suggest a gradual increase in the cytokine levels. Resultantly, we show transition in PR following an all-or-none switch in response to a graded decreasing stimulus(Fig. 8d). Such a process stresses that it occurs through as a sudden or abrupt loss of tolerance limit in the specific physiology. The timing of such a jerk to the system is varied, where, as mentioned earlier, *f_crit_* can appear at different times to different patients.

## 5 Discussion

Palindromic rheumatism has been the subject of numerous studies, yielding mixed results regarding its potential progression to rheumatoid arthritis. Some research supports the notion that palindromic rheumatism may act as a precursor to rheumatoid arthritis, indicating a possible continuity between the two conditions. Conversely, other studies dispute this connection, suggesting that it is based on RF/MEFV mutations [19]. One of the longest longitudinal studies (1967-1984) [15] reported a 63% turnover of PR patients developing chronic RA, over a period of 10 years. Joints like the proximal interphalangeal (ICP), metacarpophalangeal (MCPs), and wrists are predominantly affected, with a joint-attack distribution akin to that observed in RA [1]. On the contrary, an 11-year follow-up revealed that there was no discernible difference in the development of RA between patients who tested positive or negative for ACPA (Antibodies to Citrullinated Proteins), disproving the argument for the presence of the risk factor [12][11]. Clinical ultrasound imaging reveals the presence of inflammation occurring exterior to the joint capsule without simultaneous synovitis in individuals with palindromic arthritis (PR). This stands in contrast to RA, where inflammation within the synovial membrane is a defining feature [10]. Hence, numerous conflicting studies concerning the PR-RA transitive argument highlight gaps in understanding the immune repertoire response both preceding the onset and throughout the progression of the disease. For a comprehensive understanding of palindromic rheumatism, it is viewed as part of the preliminary phase of RA with a solely relapsing-remitting manifestation of the full disease, with a slightly less-aggressive inflammatory signature.

In the Baker Model, the palindromic state is described as an intermediary phase between the physiological and sustaining RA states [7]. However, our modified model defines PR as a separate identity, where patients may remain PR or transit to chronic disease, driven by an embedded biological mechanism. The model is capable of generating transient flares, that represent synovitis in clinical settings. The flares constitute of robust biological properties within the bistable-flare region. Our understanding on the separate fluxes suggest that it’s a self-regulatory process, where the positive feedback is effectively controlled by delayed negative feedback from the *a* → *p* interaction. Interestingly, the subtlety of such acute inflammatory events is the subsequent decay to the same equilibrium levels from where it originated. It is well recognized that flare-ups of RA can be disabling and exacerbate inflammatory conditions [20]. Thus, intuitively, we show that synovial flares in early RA are likely to rest in a slightly higher equilibrium (Fig 5), even if the change is miniscule along the disease continuum. Our attempt clearly contrasts palindromic patients that can sustain the disease’s syndromic condition on its own. A particular study in sero-negative PR patients showed upregulation of IL10RA, CXCL16 and apoptotic pathways of regulation [21]. Other studies speak of abrupt synovial flares in palindromic patients [10], we showed such flares depending on only the abrupt presence of triggers in the system. PR patients remain intermittent for many years [8], before attaining clinical persistence. Instinctively, it suggests that the synovial flares may remain as an independent recurring event, loosely connected with systemic inflammation for the time-period. There is ongoing discussion on whether PR should be viewed as a component of the spectrum of RA or as a separate illness with similar risk factors [10].

These short-term, excitable changes in the physiology, however, point to a primed autoimmune system that can self-enable disease, but in what situations? In order to answer this question, we added a regulatory control on the linear influx of *P* where the effect of *A* was found to be weak. Our mathematical argument stand on establishing strong negative regulation onto the influx of cytokines. We argue that the palindromic state would have modulatory inhibitory coupling with the system, and will not be incidental on the concentration of *A* in the system. Consequently, it implies that the system would be severely constrained during intermittent rest times in PR patients, when physiological cytokine levels are low.

In biological systems, even when the signal is present for an extended period of time, an adaptive response to an external stimulus frequently returns to its pre-stimulus value [22]. Our modified model described PR as a adjustable bistable system, where all scales of disturbances were returned to pre-stimulation status. No matter how many incapacitating triggers the PR system is exposed to, it will rest at physiological levels of cytokines in between flares. However, continued insults lead to aberration signals in the system. In cardiac physiology, the right ventricle frequently adapts to high pressure by becoming hypertrophic. This adaptation, however, frequently turns maladaptive in diseases like diabetes and pulmonary vascular disorders, resulting in a decreased ejection fraction and cardiac dysfunctionality [23, 24]. Similar principles survive in the progression of chronic kidney disease and metabolic roles of adipose tissue [25, 26]. A review article in major depressive disorder (MDD) stated chronic stress leads to innate immune activation, producing pro-inflammatory cytokines. However, sustained stress levels desensitizes T-cell inhibitory effects on cortisol, leading to mal-adaptiveness with a decreased activation of T-reg cells [27]. Comparably, we showed that a secondary adaptation brought on by repeated attacks over time may eventually turn maladaptive, which could explain the change observed in PR patients as a breakdown of tolerance.

In rheumatoid arthritis (RA), several studies have assayed the levels of cytokines to explore the ideation of bioclinical markers [28, 29, 30]. Since the inhibitory effects of anti-cytokines are vaguely understood, there is a dire need for clever research analysis to identify the molecular mechanisms that define *f*. This opens up numerous experimental avenues that remain largely unexplored. Numerous studies show how palindromic patients develop chronic disease [1, 8]. Our model offers a conceptual foundation for comprehending the shift to RA in a qualitative manner. The major limitation of our study is the need for more validation of the significant processes described for the recovery of metastable insults, as well as digression to chronicity. While accepting that there are multiple subtypes of PR [10], our study proposes transition in the subset of PR patients with a higher odds of transition to RA. To identify the mechanisms and cellular processes that might be causing this transformation, a great deal of research is required to look at changes in the genome, epigenome, and translation in longitudinal animal studies.

## 6 Appendix

### 6.1 Flux analysis

We show the individual flux rates involved in the evolution of pro– and anti-inflammatory cytokines, during a synovial flare, capturing the essential dynamics that are necessary to explain the rise and fall of cytokines to a baseline state. There are fours phases into which the flux rates are divided (Fig. 9). The biological description in each of these phases is described below.

**Figure 9:**
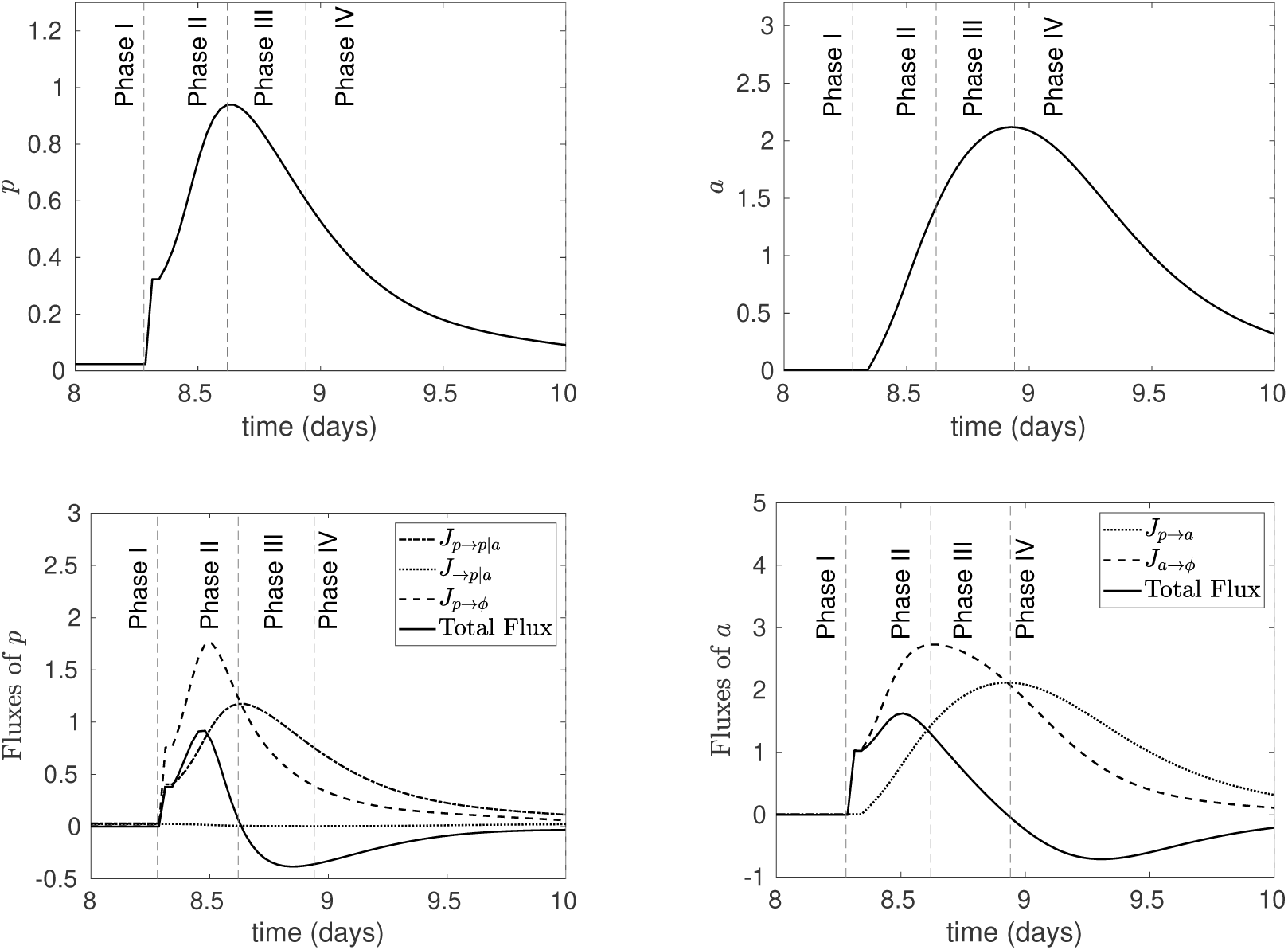
The above separate fluxes are plotted against time, with aligned phases. (a) Flare trajectory of *P* (b) Flare trajectory of *A* (c) Individual rate fluxes of *P* (d) Individual rate fluxes of *A*. The rate for the auto-feedback flux occurs at the shortest timescale, while the degradation fluxes in both variables follow a longer timescale. (The legend labels and parameters are consistent with Section 2)

**Figure 10:**
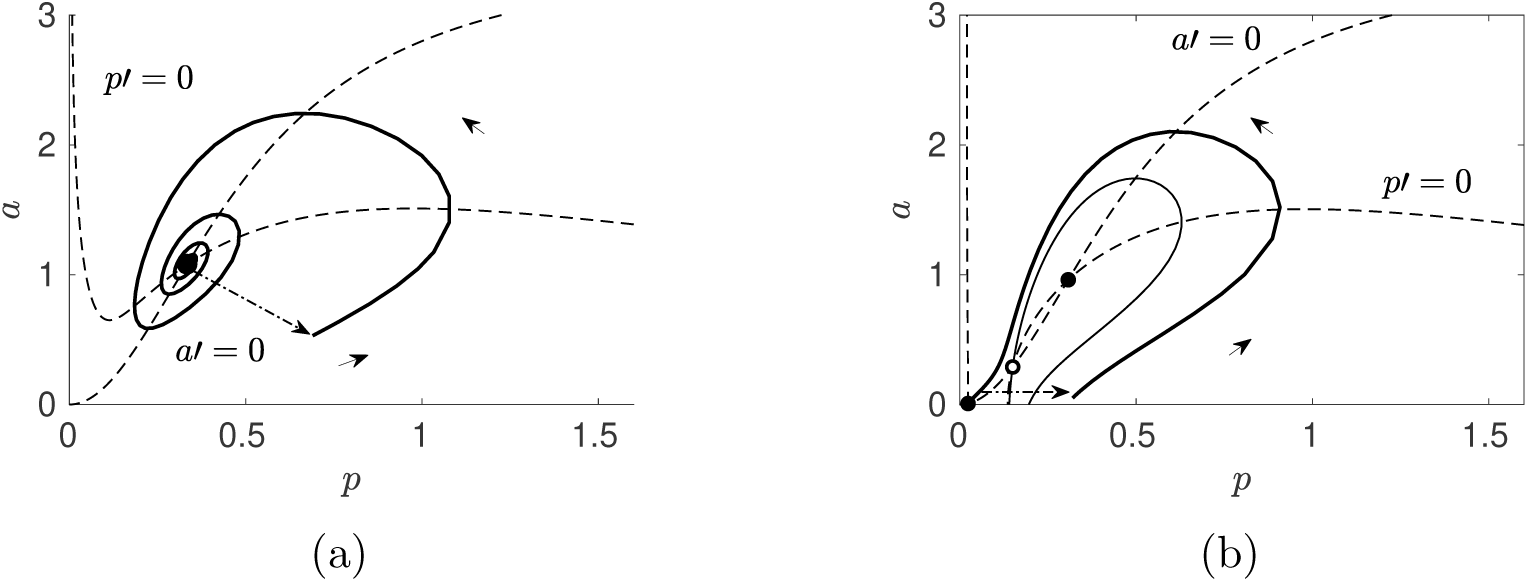
The stable steady states (s.s.) are represented by closed circles, and unstable/saddle nodes are represented by open circles. (A) Flare M has a single stable steady state. (*α*_1_=0.1) (B) Flare B has two stable steady states, separated by an unstable steady state. (*α*_1_=0.025) The other parameters are as described in Section 2.

**Figure 11:**
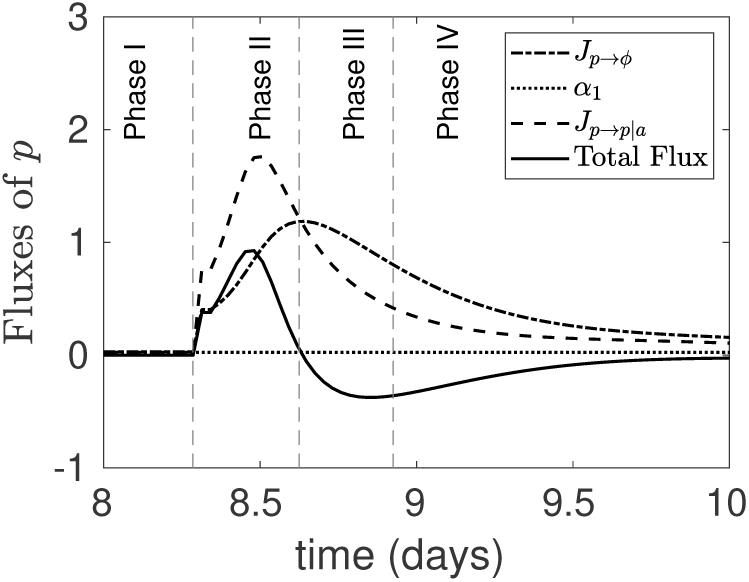
The system shows the property of flaring, in the absence of inhibition on the background flux, thus indicating that the inhibition onto *α*_1_ is not mandatory to preserve the dynamics of the system. (Parameters: Section 2)

**Figure 12:**
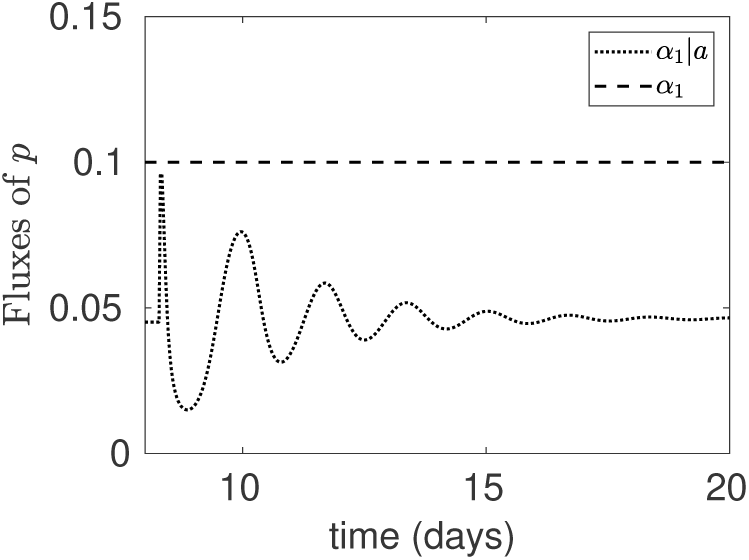
The inhibitory flux settles at a lower value, when inhibited by *a* at *α*_1_ = 0.1.

**Figure 13:**
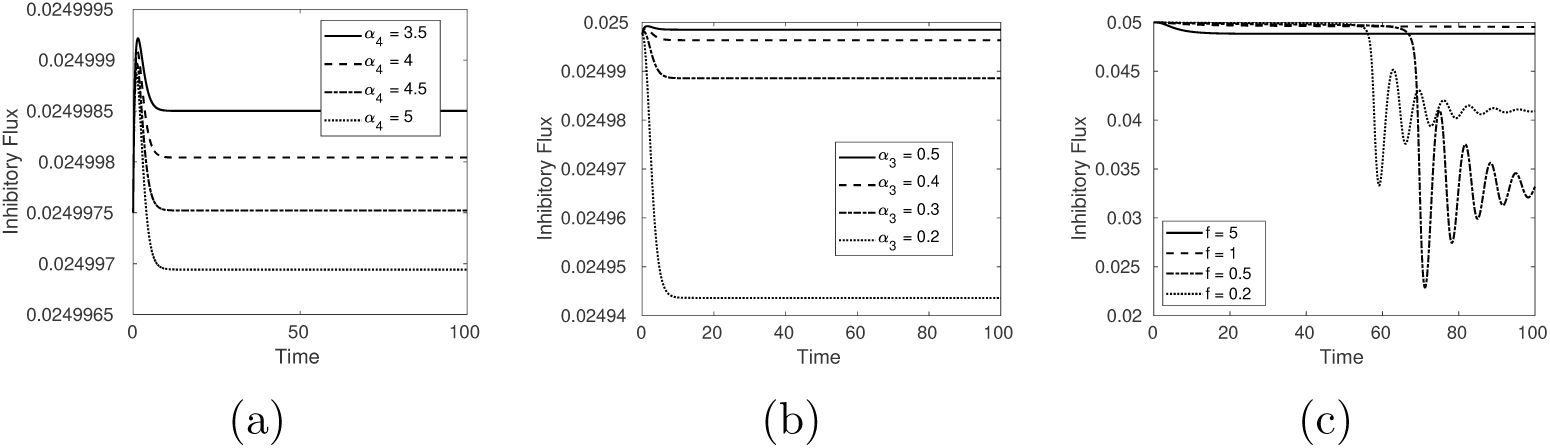
(a) We see a decrease in the inhibitory flux, with increase in *α*_4_ (b) We see a decrease in the inhibitory flux, with increase in *α*_3_ (c) *Significant* changes are observed in the inhibitory flux, with decrease in *f* (Other parameters are in Section 2)

#### Phase I

The first phase explains the unperturbed steady-state attractor positions of the *P*-*A* variables. The positive flux rates for *P*, that is, feedback flux and inhibition flux, while the negative flux rate, that is, degradation flux, do not change with time and are equally balanced, which sustains equilibrium concentrations in the *P* trajectory. Similarly, the positive flux rate for *A* i.e. production flux, negates the degradation flux, enabling sustenance at that particularly low rate as indicated in the total flux rate. Cytokine levels are undisturbed and continue to remain low.

#### Phase II

The initial trigger enables a shoot in the pro-cytokine (*P*) levels, further pushing the auto-feedback loop to take over the system. The degradation rate flux for P also undergoes a consequent increase. The delay in the rise in the degradation of P establishes the faster rise and slower fall. The resultant total flux peak halts. Once the degradation rate flux surpasses the auto-feedback flux rate, the total flux rate heads towards a steep decline, and becomes zero at the end of the phase. The rise in *P* leads to a direct delayed production in *A*, as steady state values of *A* depend on presence of *P*. With the rise in the production flux of *A*, a gradual increase in the degradation rate flux is observed, which enables a gradual decrease in the total flux rate. The dynamics of the inhibitory rate flux, although changes, but is not significant enough. The flux itself is important to reach the same steady state levels, but the nature of the stability does not change in it’s absence. The end of this phase, marks the equilibrium value obtained by the positive and negative fluxes of *P*.

#### Phase III

The auto-feedback flux decreases steeply compared to the degradation flux of *P*, creating a negative total flux. This further leads to a decrease in the production flux of *A*. The degradation flux of *A*, has a delayed increase causing the balance of the fluxes at the end of the phase. The feedback flux does not rise to a greater amplitude, due to the delayed rise in *A*. The end of this phase, marks the equilibrium value obtained by the positive and negative fluxes of *A*.

#### Phase IV

There is a similar trend absence of dip as the negative fluxes take over the positive fluxes in the system. The fluxes become unchanging at this point and is analogous to Phase I. In both the cases, the fluxes attain balance and stability in the later stage.

### 6.2 Phase plane analysis of flares for two parameter sets

We depict the phase-plane dynamics of the two flares, differing in their steady-state levels.

Flare-B has the same phase space as shown in Fig 4. Here, flare-M requires an initial perturbation that involves a reduction in *A* and an increase in *P*, while flare-B requires only an increase in *P*. Both of these flares retain the attribute of “palindromicity”, as they return to the same steady state values, regardless of where they occur along the continuity of the disease.

### 6.3 Inhibitory role of ***A*** on external drive to ***P***

#### 6.3.1 Weak inhibition

At comparatively lower values of *α*_1_, the influence of anti-inflammatory cytokines on flux remains unimportant when the system is at lower levels of *P* /*A*. The inflow of the total amount of cytokines into the motif is determined by the strength of *α*_1_ alone. If we modify the equations such that:

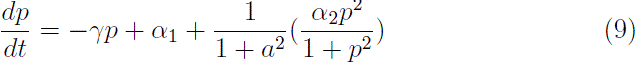

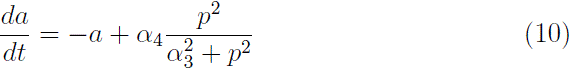

We show the individual fluxes in this case.

We considered the inhibitory influence of *A* on *α*_1_ as a constant, effectively eliminating it’s role. The dynamics of the fluxes remain the same, due to very minor changes in the steady state. Extended study of the fluxes led us to conclude that the role of the inhibitory flux were to decrease the peak-height of the auto-feedback flux.

#### 6.3.2 Strong inhibition

At comparatively higher values of *α*_1_, the influence of anti-cytokines onto the flux becomes important, when the system stabilizes at higher levels of *P* /*A*.

### 6.4 Influence of ***α*_3_**-***α*_4_** on inhibition

It becomes imperative to question the effect of the two base parameters in the model, which directly control the concentration of *A*. The effect of *α*_4_/*α*_3_ is not profound on the inhibitory flux in the previous model. However, in the modified one, the influence of *f* is significant. Here, we recall that although the parameters of the effect of *A* on *P* are taken into consideration, nondimensionalisation is not effective to account for the inhibitory effect.

## Footnotes

1 In future iterations of the model we hope to address this more mechanistically.

